# Cancer type-specific prioritization strategy for targetable dependencies developed from the DepMap profile of head and neck cancer

**DOI:** 10.1101/2022.06.12.495566

**Authors:** Austin C. Cao, Malay K. Sannigrahi, Pavithra Rajagopalan, Robert M. Brody, Lovely Raghav, Phyllis A. Gimotty, Devraj Basu

**Author notes:** **Corresponding author**: Devraj Basu, MD, PhD, 3400 Spruce Street, 5 Ravdin/Silverstein, Philadelphia, PA 19104.

## Abstract

**Objectives:** The DepMap genome-wide loss of function CRISPR screens offer new insight into gene dependencies in HPV(-) head and neck squamous cell carcinoma (HNSCC) cell lines. We aimed to leverage this data to guide preclinical studies by cataloging novel targetable dependencies that are predicted to offer a useful therapeutic window. We also aimed to identify targets potentially representing synthetic lethalities by testing for associations between genetic alterations and gene dependency profile.

**Methods:** DepMap was queried for gene probability and effect scores in cell lines from 87 tumors, including 63 HPV(-) HNSCCs plus 24 esophageal squamous cell carcinomas (ESCCs), which have comparable etiology, tissue or origin, and genetic profile to HNSCC. A probability score of ≥ 0.5 was used as the threshold for essentiality. Essential genes were selected for analysis by 4 criteria: (1) presence in ≥10% cell lines, (2) lack of common essential designation by DepMap, (3) lack of predicted dependency in normal cell lineages, and (4) designation as druggable by the Drug-Gene Interaction Database.

**Results:** The 143 genes meeting selection criteria had a median gene effect score of 0.56. Selection criteria captured targets of standard therapeutic agents of HNSCC including *TYMS* (5-FU), tubulin genes (taxanes), *EGFR* (cetuximab), plus additional known oncogenes like *PIK3CA* and *ERBB3*. Functional classification analysis showed enrichment of tyrosine kinases, serine/threonine kinases, RNA-binding proteins, and mitochondrial carriers. 90% of the 143 dependencies were not known oncogenes in the OncoKB Database. 10% of targets had inhibitors previously used in a non-HNSCC phase II trial, including 8 that have not yet been tested in cancer. The 13 genes with median gene effect scores greater than of PIK3CA and not well-studied in HNSCC were assigned highest priority, including *DHRSX, MBTPS1, TDP2, FARS2, TMX2, RAB35, CFLAR, GPX4, SLC2A1, TP63, PKN2, MAP3K11,* and *TIPARP*. A novel association was found between *NOTCH1* mutation and increased *TAP1* dependency.

**Conclusions:** The DepMap CRISPR screens capture well-studied targets in HNSCC as well as numerous genes without known roles in HNSCC or malignancy in general. Several of these targets have well-developed inhibitors that provide resources to guide preclinical studies. Association of some of the dependencies with known molecular subgroups in HNSCC may enhance use of cell line models to guide personalization of therapy.

## INTRODUCTION

Advanced HPV-negative head and neck squamous cell carcinoma (HPV(-) HNSCC) continues to carry a poor prognosis compared to its HPV-positive counterpart^1^. The emergence of modern targeted therapies such as anti-EGFR and anti-PD(L)1 agents has had limited overall impact on oncologic outcomes for these patients^2^. Other recently proposed strategies, including PI3K and CDK 4/6 inhibition, have yet to show utility in clinical trials for monotherapy or combination therapy, as they may be hampered by prognostic associations with genomic subgroups of HPV(-) HNSCC. Though subgroup classification based on genomic alterations or expression profiles is common in other cancers, literature on how biomarkers in HPV(-) HNSCC may predict immunotherapy response remains sparse. Recent efforts have focused on identifying previously unstudied driver genes, but characterization of the genetic and transcriptional landscape of HPV(-) HNSCC via the Cancer Genome Atlas and similar projects has not yet led to new targets achieving clinical application for this disease^3^.

Development of targeted therapies requires detailed knowledge of the genetic makeup of tumor cells. The DepMap Portal now offers results from pooled genome-wide CRISPR-Cas9 screens for 990 cell lines from multiple cancer types, including a large panel of HPV(-) HNSCC models^4^. CRISPR-Cas9 dropout screening offers improved identification of vulnerabilities in cancer due to its limited off-target effects. This resource may provide a new window into the molecular dependency profile for HPV(-) HNSCC that can lead to preclinical evaluation of new therapeutic approaches. While DepMap provides useful visualization tools for exploring the database, prioritization of cancer-specific dependencies remains a challenging task for researchers interested in evaluating potential drugs in cell lines. We aimed to guide use of cell line models for preclinical studies by cataloging recurring, targetable dependencies revealed by DepMap that are predicted to offer a cancer-specific therapeutic window. In addition, we aimed to identify the subset of targetable dependencies representing potential synthetic lethalities by evaluating for associations between the common genetic alterations in the cell lines and their targetable dependency profiles. Overall, we sought to propose a target prioritization strategy for cell line models of HPV(-) HNSCC that could be applied to any cancer type available in DepMap.

## METHODS

### DepMap

The DepMap Public 21Q3 CRISPR screening library contains CRISPR-Cas9 knockout data for 17,387 genes in 990 cell lines^1^. All HPV-negative head and neck squamous cell carcinoma (HNSCC) cell lines annotated in DepMap were selected for analysis (n=63). The size of this panel was increased to 87 by adding all 24 cell lines from esophageal squamous cell carcinoma (ESCC), which have closely comparable etiology, tissue of origin, and genetic landscape to HPV(-) HNSCCs^5, 6^. DepMap reports a gene probability score as the probability (0-1) that knocking out a gene has a true effect on cell line survival, and they use a score > 0.5 to designate a gene as essential to a specific cell line. To identify recurring dependencies, we used this cutoff to filter for genes essential in at least 10% (8/87) of the cell line models. DepMap also reports a gene effect score as the effect size of knocking out a gene on cell line survival, with a negative value representing a stronger dependency. For simplification, the inverse was taken of all gene effect scores in this report such that a more positive value represents a stronger dependency. To reduce false positives, scores are corrected using CERES, a computational method accounting for copy-number-specific effects and variable sgRNA activity. The scores are finally scaled such that the median gene effect score for nonessential genes across all cell lines in DepMap is 0 and the median for essential genes is 1. In our analysis, genes with the highest median gene effect scores in the 87 cell line models were assigned the highest priority for further study.

### Common essential and core fitness genes

We used two sources to exclude dependencies that are shared by normal tissue and thus would not provide a useful therapeutic window. First, DepMap designates “common essential” genes as dependencies that are present across >90% of all 990 cancer cell lines^7^. Second, we identified genes that were dependencies across 3/5 normal cell lineages in prior pooled loss-of-function CRISPR screens^8^. Common essential and core fitness genes were therefore excluded due to the increased likelihood of causing toxicity to normal tissue.

### DGIdb

Potential targetability of gene products was evaluated using the Drug Gene Interaction Database (DGIdb), which aggregates data from multiple sources for existing and potential drug-gene interactions^9^. DGIdb designates genes in the “druggable genome” if a drug-gene interaction exists or if the gene belongs in one of 42 potentially targetable gene categories, such as kinases and ion channels. To best identify targetable dependencies, genes not part of the “druggable genome” were excluded from analysis.

### Open Targets

To characterize the status of drug development for genes of interest, the list was crossed with the Open Targets Platform, which integrates publicly available datasets to curate approved or investigational drugs known to directly act on gene products^10^. Clinical trial phase and disease indications were collected for genes selected in our analysis. Gene targets with an “existing clinical agent” were defined as having an inhibitor studied in a previous or current Phase II trial for any disease indication, which indicates the presence of an established safety profile. Genes with an existing clinical agent in head and neck cancer were identified in the prioritization pipeline.

### OncoKB

The OncoKB database annotates genes as oncogenes or tumor suppressor genes based on their inclusion in various sequencing panels, the Sanger Cancer Gene Census, or Vogelstein et. al^11–13^. OncoKB is the most inclusive database of cancer-related genes. To validate the filtration steps in this pipeline, OncoKB was used to screen the gene list for well-studied targets in cancer.

### DAVID Gene Functional Classification Tool

DAVID Bioinformatics Resources is an open database that integrates biological data and analytical tools for functional annotation of genes and pathways^14^. The DAVID Gene Functional Classification Tool was used to classify gene lists into functionally related groups based off 14 functional annotation sources, including gene ontology terms (GO)^15^. Significant enrichment of a group within our gene list was defined using a kappa similarity threshold of 0.3 and similarity term overlap of 4. We therefore used DAVID to summarize recurring functional classifications in our pipeline.

### Identification and prioritization of targetable dependencies

We describe the prioritization pipeline for targetable dependencies in HPV(-) HNSCC cell line models utilizing the data sources listed above. DepMap was queried for gene probability and effect scores in cell lines from 87 tumors, including 63 HPV(-) HNSCCs and 24 ESCCs. A gene probability score of ≥ 0.5 was used as the threshold for essentiality for a specific cell line. Targetable dependencies were identified using four criteria: (1) Essential in ≥ 10% cell lines, (2) lack of common gene essential designation by DepMap, (3) lack of dependency in CRISPR screens of normal human cell lineages, and (4) designation as potentially targetable by DGIdb. Targetable dependencies were further prioritized for consideration of further study using these criteria: (1) Lack of an existing clinical agent in HNSCC, (2) median gene effect score greater than that of PIK3CA (0.57), which is a well-studied oncogene in HNSCC with the weakest dependency in our analysis.

### Associations between common genetic alterations and targetable dependencies

To identify common genetic alterations which may confer unique vulnerabilities in cell line models, we aggregated a list of genes with frequent and significant genetic alterations in HPV(-) HNSCC from two review articles^3, 16^. The Cancer Cell Line Encyclopedia (CCLE) includes a detailed genetic characterization of human cancer models, including all 87 cell lines used in this analysis^17^. The CCLE was queried for the presence of relevant alterations, depending on their classification as an oncogene or tumor suppressor in the review articles. Genes with alterations in ≥ 10% cell lines in our pipeline were included in the associations analysis. Associations between common genetic alterations and targetable dependencies were defined using two-sample t-tests comparing the median gene effect score between cell lines with the genetic alteration and without the genetic alteration^18^. Significant associations were defined using filter conditions of p < 0.05 and Cohen’s d effect size ≥ 1.

## RESULTS

### Identification of targetable dependencies in cell line models of HNSCC

All HPV-negative head and neck squamous cell carcinoma (HNSCC) cell lines annotated in DepMap were selected for analysis (n=63). The size of this panel was increased to 87 by adding all 24 cell lines from esophageal squamous cell carcinoma (ESCC), which have closely comparable etiology, tissue of origin, and genetic landscape to HPV(-) HNSCCs^6, 19^. Starting with the 17,387 genes targeted in across the DepMap CRISPR screens, the gene probability score cutoff used by DepMap to designate genes as essential to survival (>0.5)^20^ indicated 5123 genes to be essential in at least one of the 87 cell lines **(Figure 1)**. From these 5123 genes, a subset of 1001 was selected based on prediction by the Drug-Gene Interaction Database that their protein products are therapeutically targetable.^9^ The 1001 genes were visualized as points on a scatter plot showing the percentage of cell lines that consider a gene essential (x-axis) and the median gene effect score in those cell lines (y-axis) (**Figure 1A**). A high frequency of genes is clustered at the low and high ends of the y-axis. This analysis provided a large list of potentially targetable dependencies in HNSCC cell lines that are listed in **Supplementary Table 1**.

**Figure 1:**
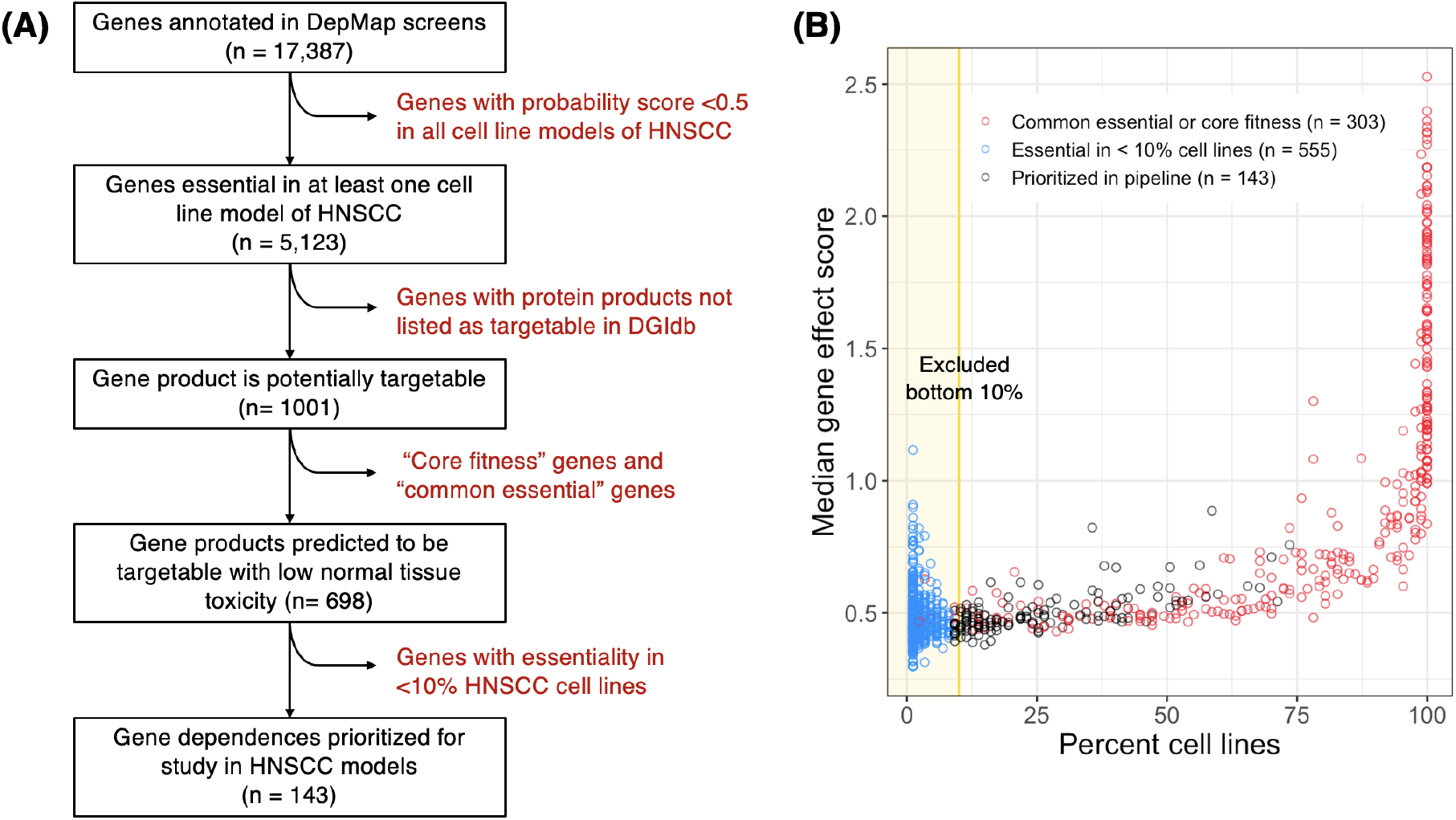
Identification of targetable dependencies in HNSCC. (A) Schematic overview, and (B) Percentage of cell lines with essentiality compared with median gene effect of all targetable dependencies (n = 1001).

To prioritize among the 1001 targetable dependencies, we considered the following features: (1) the percentage of cell lines that consider a gene to be essential (2) the median strength of dependency in those cell lines, and (3) whether the dependency is shared by nonmalignant cells. First, we sought to remove gene dependencies that are shared by normal tissue and thus would not provide a useful therapeutic window. The 1924 dependencies that are present across >90% of all 990 cancer cell lines in DepMap have been designated as “common essential” genes, which are predicted to lack specificity to cancer, and 160 such common essential genes were present in the list of 1001 targetable dependencies for HNSCC. These genes had a median gene effect score of 1.28 (IQR: 1.00-1.79), which is consistent with the scaling of DepMap gene effect scores to set 1.0 as the median for all common essential dependencies. To further assess for dependencies shared by normal tissue, we identified genes that were dependencies across multiple normal cell lineages in prior pooled loss-of-function CRISPR screens [2015]. Of the 1580 such “core fitness” genes designated in this prior study, 190 were present in the gene list of targetable dependencies. The 190 core fitness genes had a median gene effect score of 0.92 (IQR: 0.61-1.44), and 114 of them also appeared on the list of common essential genes. Presence of this large overlap provided cross-validation of these two independent approaches for excluding targets likely to have high toxicity. Both groups together comprised 303 unique genes, which are labelled in **Figure 1A**. The median gene effect scores of these dependencies are shown to be significantly stronger than that of the remaining gene dependencies (**Figure 1B**). Next, we considered the rarest gene dependencies, which comprised a large cluster of genes (n=555) appearing in ≤ 10% of the cell line panel (**Figure 1A**). These gene dependencies had significantly weaker median effect scores than others in the pipeline (**Figure 1B**), and outliers in the group with higher median effect scores were interpreted as likely arising from stochastic effects due to the small sample sizes (< 8 cell lines). The relative weakness of these 587 gene dependencies plus their rarity in the cell line panel led us to exclude them. Excluding rare dependencies along with the targets with high likelihood of causing toxicity left 143 genes that were prioritized for further analysis, and the complete pipeline leading to this gene list is summarized in **Figure 2**. These 143 genes are visualized in **Figure 3**, where a moderately strong positive relationship was apparent between frequency of essentiality and median gene effect score (Spearman’s ρ = 0.63). The 143 genes are described in **Supplementary Table 2**, where they are ranked primarily by frequency of essentiality and secondarily by median gene effect score.

**Figure 2:**
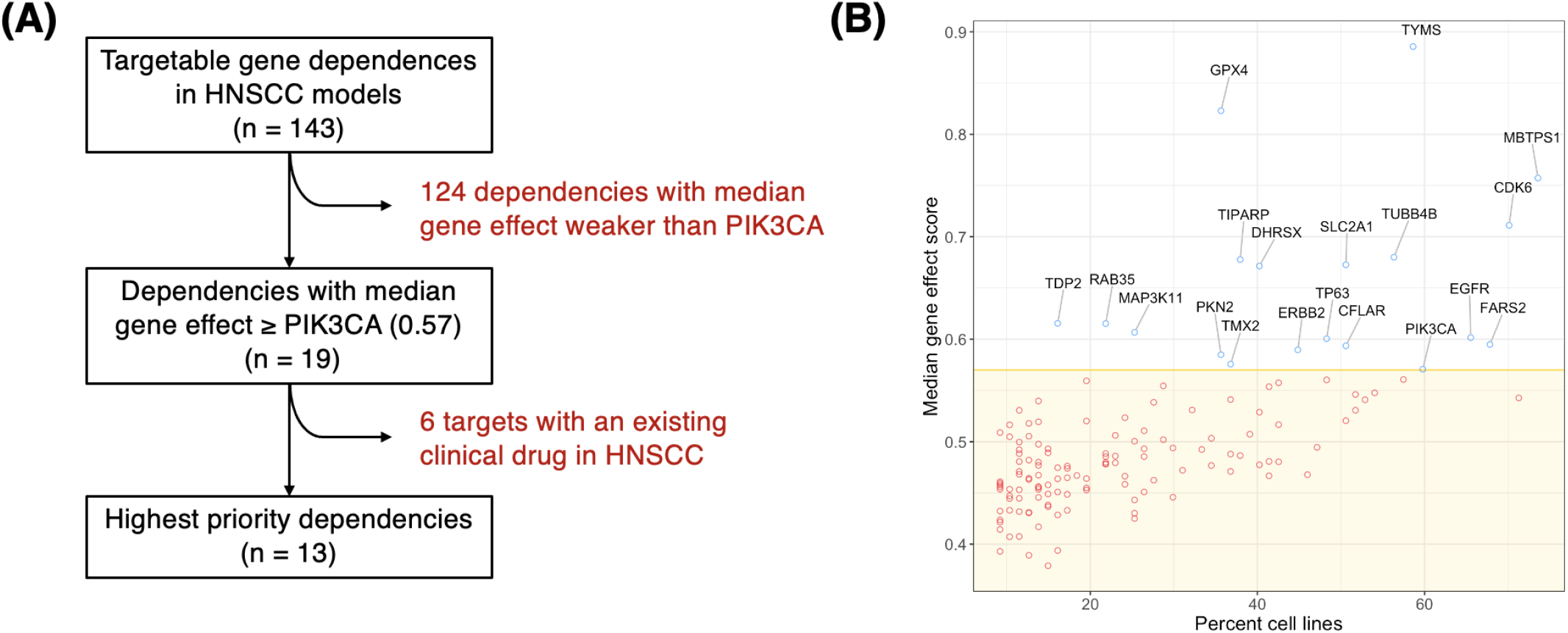
Prioritized dependencies with highest median gene effect scores: (A) Schematic overview, and (B) List of genes (n = 19) with median gene effect greater than that of PIK3CA (0.57).

**Figure 3:**
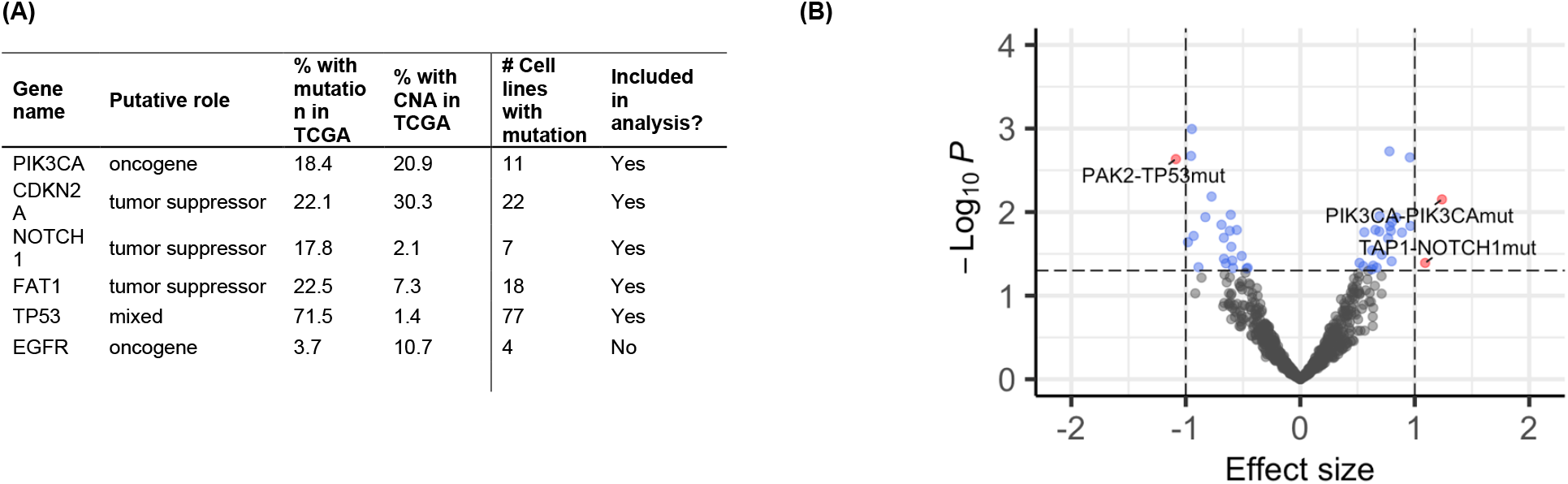
Association between median gene effect score and mutations in cell line models of HNSCC: (A) Known mutations in HNSCC found in > 10% of cell lines, and (B) Comparison with median gene effect score, with significant associations highlighted in red. A positive effect size represents increased dependency when the mutation is present.

We analyzed this list of targetable dependencies for presence of known oncogenes designated in the OncoKB database. Overall, 17% (24/143) of the genes appeared in the OncoKB database of cancer-related genes, where 10% (14/143) were designated as well-established oncogenes: *EGFR, PIK3CA, ERBB2, ERBB3, CDK6, TP63, IGF1R, RAB35, KLF5, FGF19, CDK8, PTPN1, PRKACA,* and *RPS6KA4*. In addition, the pipeline captured the molecular targets of cytotoxic drugs used in HNSCC, including thymidylate synthetase (target of 5-FU) and tubulin (target of taxanes).^21^ Despite the limitations of in vitro CRISPR loss-of-function screening, many well-studied targets in head and neck cancer were readily identifiable in the proposed pipeline.

### Functional classification of targetable dependencies

To further classify the dependencies into functionally relevant groups that may not be well-studied in cancer, the Gene Functional Classification Tool from DAVID Bioinformatics Resources was utilized. This analysis captured five enriched clusters based on similarities in their functional annotations, representing serine/threonine kinases, tyrosine kinases, mitochondrial carriers, RNA splicing proteins, and transmembrane proteins, shown in **Table 1**. Serine/threonine kinases had the strongest enrichment score and were well represented in our pipeline. A subgroup of these serine/threonine kinases including *MAP3K11*/MLK3, *MAP4K2/GCK,* and *RPS6KA4/MSK2* are involved in the ERK/MAPK signaling pathway, which is an emerging target in HNSCC^22^. Furthermore, *PKN2* and *PAK2* are two other serine-threonine kinases with strong dependencies that have not been previously evaluated as driver genes in HNSCC. Tyrosine kinases are the second most enriched group in the gene list and include well-studied targets: EGFR, *ERBB2, ERBB3, TYRO3,* and *IGF1R*. Mitochondrial transporters are also enriched, but demonstrated relatively weak dependencies and were only essential to a mean 12% of cell lines. Lastly, a notable group of RNA splicing proteins were found in the list of targetable dependencies, including *HNRNPA1, ELAVL1, ZRANB2, RBM5, RBM10,* and the well-studied *PTBP1*. Altered regulation of RNA splicing is an emerging driver of cancer, and preclinical studies on pharmacologic modulation have focused on exploiting vulnerabilities in cancer cells with altered splicing machinery^23^. Aside from *PTBP1,* the role of these specific dependencies in HNSCC is unknown and warrants further investigation. In addition to well-studied targets in head and neck cancer, recurring functional classifications with emerging roles in cancer were found among the targetable dependencies generated in our pipeline.

**Table 1.** Functional groups enriched among the 143 prioritized dependencies.

| Functional classification | Enrichment score | Genes (Median gene effect score) | Mean of median gene effect scores | Mean of % cell lines with essentiality |
| --- | --- | --- | --- | --- |
| Serine/threonine kinase | 3.24 | MAP3K11 (0.61), PKN2 (0.58), MARK2 (0.55), PAK2 (0.52), CDK8 (0.46), MAP4K2 (0.45), RPS6KA4 (0.42) | 0.51 | 22% |
| Tyrosine kinase | 2.93 | EGFR (0.60), ERBB2 (0.59), ERBB3 (0.55), IGF1R (0.54), TYRO3 (0.49) | 0.55 | 38% |
| Mitochondrial carrier | 1.35 | SLC25A33 (0.47), SLC25A1 (0.44), MTCH2 (0.41), SLC25A25 (0.39) | 0.43 | 12% |
| RNA-binding protein | 1.26 | RBM10 (0.54), PTBP1 (0.53), ELAVL1 (0.49), HNRNPA1 (0.47), ZRANB2 (0.47), RBM5 (0.47) | 0.49 | 35% |
| Transmembrane protein | 1.56 | SSR2 (0.52), TM2D1 (0.52), ODR4 (0.51), KCNK7 (0.47) | 0.5 | 30% |

### Pharmacologic approaches to target the prioritized dependencies

We aimed to characterize the current status of drug development for the 143 dependencies in the prioritized list. Towards this end, we used the Open Targets database to find the most advanced status of an inhibitor in HNSCC and other diseases. Notably, 12 of the dependencies in our gene list have inhibitors previously studied in a HNSCC phase II trial. These were generally stronger and recurring dependencies, with a mean gene effect score of 0.72 and frequency of dependency in cell lines of 47%. Seven of these dependencies have an inhibitor in a phase II/III trial but have yet to achieve approval. To explore the possibility of repurposing existing non-HNSCC agents for use in HNSCC cell lines, we also sought to identify dependencies with an existing “clinical inhibitor,” which are drugs that have reached a phase II clinical trial in other cancers or non-malignant diseases. **Table 2** shows the current status of drug development for 14 dependencies with clinical inhibitors that have yet to be used clinically in HNSCC, including 8 with FDA approved inhibitors: *EGLN1, P2RY6, PCSK9, UGCG, IMPDH1, HCRTR1, GANAB, TNFRSF8/CD30*. Overall, 7 genes have been previously targeted in a phase II cancer trial, and they may be more readily repurposed for trials in HNSCC due to known safety and efficacy in cancer: *PCSK9, IMPDH1, TNFRSF8/CD30, BIRC2, ITGB1, MAP3K11* and *LDHA*. Of these, the integrin subunit *ITGB1 was* essential to the greatest percentage of cell lines (48%) and had the strongest median gene effect score (0.56). While inhibitors targeting integrin adhesion receptors have not provided clinical utility in trials for other cancers, this data suggests that there may exist a sub-population in HPV-negative head and neck cancer that may benefit if appropriately identified. Of the genes targeted only in noncancer trials, *MAP3K11* demonstrated the most robust dependency (0.61). *MAP3K11* encodes the serine/threonine kinase MLK3, which has recently been a promising target in preclinical studies of breast and ovarian cancer^24^. These studies used CEP-1347, a small molecule inhibitor of MLK3 whose safety in humans was demonstrated in a large-scale phase III clinical trial in Parkinson’s disease. In conclusion, a small cohort of dependencies in cell line models have been clinically targeted in non-HNSCC diseases, demonstrating a safety profile in humans and making them promising candidates for repurposed inhibitor studies.

**Table 2.**
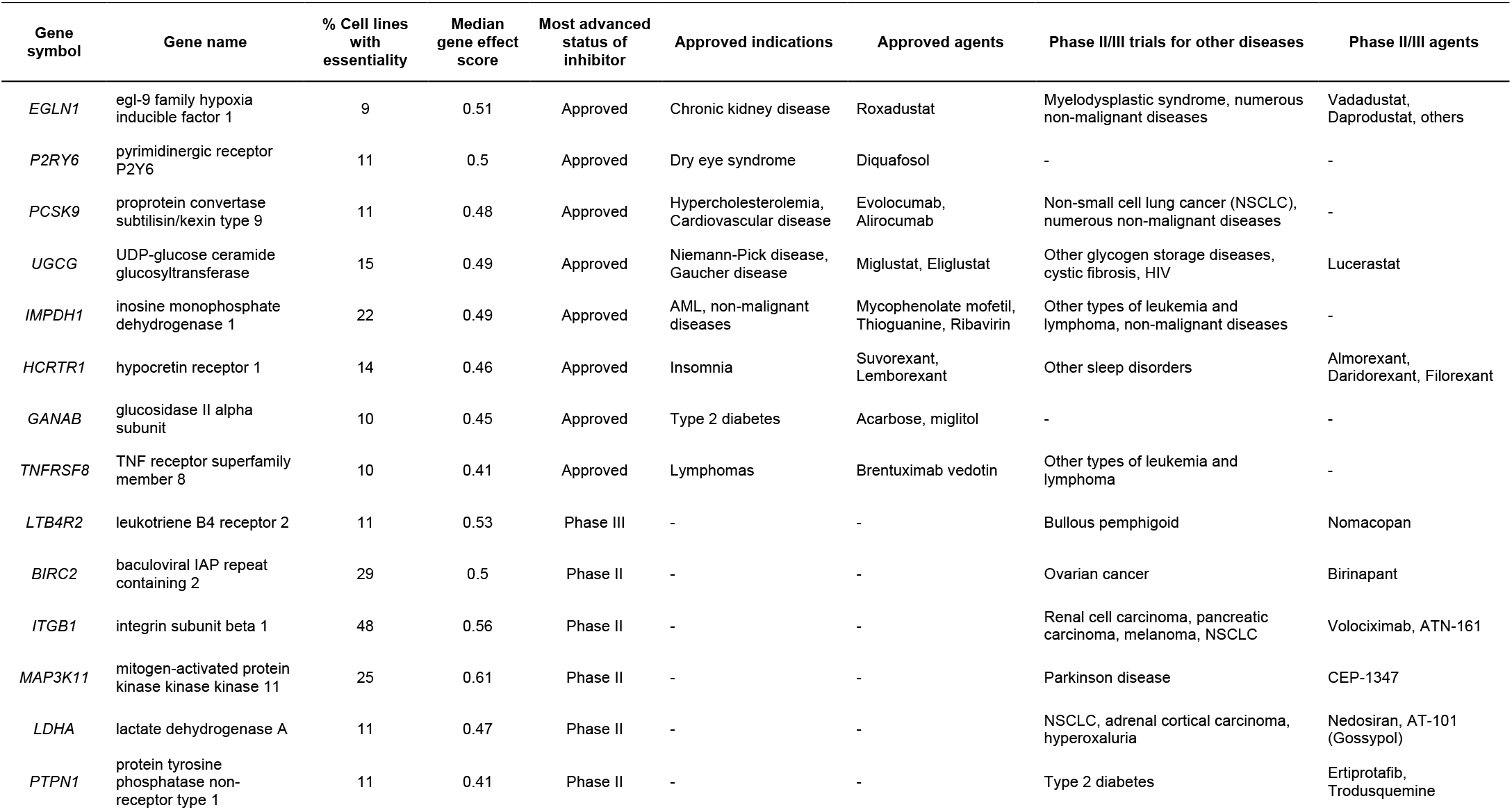
Prioritized gene products with clinical inhibitors not well-studied for HNSCC. “Clinical inhibitor” defined as an inhibitor that reached phase II trial for any disease indication.

### Identifying prioritized dependencies with highest median gene effect scores

We next sought to find targetable dependencies with the highest median gene effect scores and most likely to be a useful candidate for future studies. To establish a meaningful cutoff for median gene effect score, we referenced *Leemans et al.* for well-established and targetable oncogenes in HPV(-) HNSCC, which included *EGFR* and *PIK3CA*^25^. *PIK3CA* was found to have the lower median gene effect score (0.57), which we used as a threshold for assigning highest priority to genes. While an inhibitor for *PIK3CA* has yet to reach FDA approval for HNSCC, it remains a well-known driver of disease, and genes with higher median gene effect scores than *PIK3CA* are more likely to be clinically useful. Overall, 19 dependencies were as strong or stronger than *PIK3CA* in our cell line panel, which are visualized on a scatter plot in **Figure 3B**. To find novel targets, we next excluded the 6 genes that already have a clinical inhibitor in HNSCC, which are described separately in **Supplemental Table 2**. This addition to the prioritization pipeline is depicted in **Figure 3A** and captured a list of 13 genes most likely to be clinically valuable. Two of these dependencies *FARS2* and *DHRSX* have not been previously investigated in cancer. Other dependencies were found to be implicated in various hallmarks of cancer^26^. *CFLAR* (CASP8 and FADD like apoptosis regulator) and *TDP2* (tyrosyl-DNA phosphodiesterase 2) are involved in resisting cell death^27, 28^. *TP63* (Tumor Protein 63), *MAP3K11* (mitogen-activated protein kinase kinase kinase 11), *PKN2* (protein kinase N2), *Rab35* (member RAS oncogene family) and *TIPARP* (TCDD-inducible poly-ADP-ribose polymerase) are involved in sustaining proliferative signaling, invasion and metastasis^29–33^. *SLC2A1* (solute carrier family 2 member 1), *GPX4* (glutathione peroxidase 4), *TMX2* (thioredoxin related transmembrane protein 2) and *MBTPS1* (membrane bound transcription factor peptidase, site 1) are involved in regulating cellular energetics^34–37^. These findings support the capacity of our filtration pipeline to enrich for therapeutic targets with potential roles in cancer based on median gene effect score. These targetable dependencies should be prioritized for further investigation.

### Associations between common genetic alterations and targetable gene dependencies

Genetic alterations known to cause cancer may confer unique vulnerabilities which may be compelling therapeutic targets. To this end, we aimed to discover associations between druggable dependencies and known molecular alterations in head and neck cancer. The purpose of this analysis was to identify dependencies in subgroups of HNSCC with known driver mutations and copy number alterations. To identify common genetic alterations, we aggregated a list of genes with frequent and significant alterations in HPV(-) HNSCC from two review articles.^13,14^ The CCLE was queried for the presence of these alterations, depending on their classification as an oncogene or tumor suppressor in the review articles. Genes with alterations in ≥ 10% cell lines in our pipeline were used in the associations analysis: *PIK3CA, CDKN2A, NOTCH1, FAT1, TP53,* and *EGFR*. The inclusion of mutations or copy number alterations is shown in **Figure 3A**, and significant associations between a mutation and a dependency are highlighted in **Figure 3B**.

Mutations and copy number alterations were available for only 48 cell-lines. *FAT1* deletion is over-represented in their association with respective gene dependency. *FAT1* is frequently mutated in cancer and is known to activate various signaling pathways through protein-protein interactions and is involved in sustaining proliferative signaling, activating invasion and metastasis^38^. Recent studies show that *FAT1* over-expression suppressed the migration and invasion capability of HNSCC cells and non-synonymous *FAT1* mutations were associated with poor disease-free survival in HNSCC patients^39^. The genes associated with *FAT1* include *MBTPS1, TMX2, ADAMTS7, ARTN, CSF3, IMPDH1, ZRANB2* and *ADAM1* are novel and warrant further investigation.

Mutation data was available for 87 cell-lines. Associating them with gene dependency revealed positive self-association between *PIK3CA* mutation and *PIK3CA* gene dependency, which has been well-studied in HNSCC. The *PIK3CA* activating mutations E545K and H1047R are commonly observed in HNSCC that leads to PI3K overactivity and considered a predictive biomarker for treatment selection^40^. *TP53* mutation which results in its loss of tumor suppressor activity in HNSCC^41^ was observed to be negatively associated with its down-stream effector protein *PAK2* (p21 activated kinase 2). Wild type p53 prevents Cdc42/Rac1 dependent cell effects that control actin cytoskeletal dynamics and cell movement^42^. Cdc42/Rac1 activates PAK2 by tyrosine phosphorylation^43^. PAK2 is shown to upregulates c-Myc expression, which, in turn, transcriptionally activates and induces pyruvate kinase M2 (PKM2) expression, resulting in reduced aerobic glycolysis, proliferation, and chemotherapeutic resistance of HNSCC cells^44^. A positive novel association was also observed between *NOTCH1* mutation and *TAP1* gene which requires further investigation. Thus, association of dependency profiles with known molecular subgroups enhances the application of cell line models to the development of personalized therapeutics.

## DISCUSSION

In this study, we developed a comprehensive methodology for prioritizing cancer-specific vulnerabilities based on potential targetability, strength of dependency, frequency of dependency, lack of dependency in normal cells, and lack of prior clinical study in a specific disease. When applied to 87 models of HPV(-) HNSCC available in DepMap, our methodology identified 143 targetable dependencies, including 13 genes designated as highest yield for future preclinical inhibitor studies. We propose a cancer-specific prioritization strategy that may guide researchers seeking to identify clinically useful dependencies in other cancer types.

CRISPR screening has been leveraged as a powerful tool for understanding complex gene functions and identifying genetic dependencies with a high degree of precision. DepMap provides a large map of vulnerabilities found in cancer cell models using CRISPR loss-of-function, which may be helpful for understanding their roles in diverse biological processes and their potential as druggable targets in cancer. The sensitivity of this pipeline was demonstrated by capture of targets of standard therapeutic agents including thymidylate synthase, tubulin, and EGFR, in addition to other well-known oncogenes in HNSCC like *PIK3CA, ERBB3,* and *CDK6* among the list of 143 targetable dependencies. Targeted therapies for many of these oncogenes are already in ongoing trials for HNSCC, including *PIK3CA* inhibitors like buparlisib, alpelisib, BKM120 and BYL719 [NCT04338399, NCT04997902, NCT01816984, NCT02537223] and *CDK6* inhibitors palbociclib and abemaciclib [NCT04966481, NCT04169074, NCT03356223, NCT03356587]. Overall, 83% of the dependencies identified were not known cancer genes in the OncoKB database and may represent novel vulnerabilities in cancer cell models. Functional classification analysis revealed significant representation of proteins associated with different mechanisms in tumorigenesis suggesting potential biologic relevance, including tyrosine kinases, serine/threonine kinases, RNA-binding proteins, and mitochondrial carriers. Furthermore, proteins that have been previously targeted in other cancers or nonmalignant diseases are well-represented in our pipeline, indicating prior safe use in humans. This subset of dependencies provides readily available drugs that can be repurposed for study in HPV(-) HNSCC cell line models. Of these, the serine/threonine kinase *MAP3K11/MLK3* was the only target with a median gene effect score greater than PIK3CA, a well-studied oncogene in HNSCC and a useful benchmark for determining potential clinical usefulness. The MLK3 inhibitor CEP-1347 has been shown to be safe in a large-scale phase III trial for Parkinson’s disease and may be a promising candidate for study in HNSCC. Overall, these findings support the robustness of this prioritizing pipeline to capture well-studied targets in HNSCC and other clinically relevant targets with established safety profiles.

Identification of genes for which tumor cells shown addiction is a promising approach for discovery of molecular targeted therapy. We used the strength of addiction, represented by median gene effect score, to prioritize 13 dependencies that were as strong as the oncogene PIK3CA in our pipeline but have not been previously studied in HNSCC. Two of these dependencies, *FARS2* and *DHRSX,* have not been implicated in any cancer. *FARS2* encodes mitochondrial phenylalanyl-tRNA synthetase^45^ and *DHRSX* encodes a non-classical secretory protein shown to regulate autophagy during cellular starvation^46^.

Besides these, another 11 genes that have been studied to varying degrees in cancer were identified and may be clinically valuable in HNSCC based on our unbiased analysis. Here, we describe the known roles in cancer of these genes, in addition to the descriptions provided in **Table 3**. To begin, CFLAR is an anti-apoptotic protein often over-expressed in HNSCC^47^ and known to block necroptosis, thereby triggering resistance to anticancer agents. Its degradation induces apoptosis in many head and cancer cells^27, 48^. TDP2 is 5’-tyrosyl DNA phosphodiesterase that repairs DNA double-strand breaks, preventing apoptosis and potentially conferring chemoresistance^28^. TP63 is predominantly expressed as the ΔNp63α variant and has been shown to inhibit the proapoptotic p53-related protein p73, suppress p16/INK4A expression, and induce *EGFR* expression^49^, thereby increasing cell survival and proliferation^29^. MAP3K11 or MLK3 is a serine/threonine-protein kinase shown to regulate cell proliferation, migration, and invasion independent of EGF signaling^30^, and MLK3 inhibition has been proposed to enhance the effects of EGFR inhibition. We previously discussed MLK3 as the strongest dependency with a clinical inhibitor previously used for a non-HNSCC disease, making it an accessible target for preclinical study in HNSCC. PKN2 is another serine/threonine-protein kinase that functions as effectors of Rho GTPases and is an essential regulator of both entry into mitosis (G2/M progression) and exit from cytokinesis^50^., inhibiting PKN2 expression results in decreased colony formation, invasion and migration in HNSCC cells^31^. Rab35 is associated with the Rho family of GTPases and localized in both plasma membrane and endosomes. It is involved in vesicular membrane trafficking, actin dynamics and regulation of the PI3K pathway^32, 51^. TIPARP or PARP7 is a mono-ADP-ribosylating protein that modifies multiple transcription factors^33^ and negatively regulates microtubule stability, thereby inhibiting cancer cell growth and motility^52^. SLC2A1 or GLUT1 is a membrane protein that facilitates the basal uptake of glucose for most metabolic pathways to meet the energy demand of cells and has been observed to be upregulated in HNSCC^34, 53^. GPX4 is involved in glutathione metabolism, protects cells against oxidative damage, and inhibits ferroptosis^35, 54^. Accordingly, increased *GPX4* expression has been shown to sensitize HNSCC cancer cells to various anticancer drugs^36, 54^. TMX2 also protect cells against oxidative damage and its inhibition resulted in decreased mitochondrial respiratory reserve capacity and compensatory increased glycolytic activity^55^. Lastly, MBTPS1 or site-1 protease catalyzes the proteolytic activation of transcription factor sterol regulatory element-binding proteins^56^ and the cyclic AMP-dependent activating transcription factor 6^56^. It is involved in regulation of many metabolic pathways including glycolysis, citric acid cycle and fatty acid biosynthesis^37^.

**Table 3:** Functions of prioritized genes with highest median gene effect scores.

| Gene symbol | Protein name | Designation in OncoKB | Description | Mechanistic study in HNSCC? | % Cell lines with essentiality | Median gene effect score |
| --- | --- | --- | --- | --- | --- | --- |
| DHRSX | DHRSX | - | Oxidoreductase involved in positive regulation of autophagy. | No | 40 | 0.67 |
| MBTPS1 | S1P | - | Proprotein convertase involved in lipid biosynthesis and protein quality control. | No | 74 | 0.76 |
| TDP2 | TDP2 | - | Phosphodiesterase involved in DNA repair, prevents DNA damage in cancer cells. | No | 16 | 0.62 |
| FARS2 | FARS2 | - | Mitochondrial phenylalanyl-tRNA synthetase overexpressed in many cancer types. | No | 68 | 0.59 |
| TMX2 | TMX2 | - | Oxidoreductase in the ER involved with protein folding and redox regulation. | No | 37 | 0.58 |
| RAB35 | Rab35 | Oncogene | GTPase involved in membrane trafficking and activates PI3K-AKT signaling. | No | 22 | 0.62 |
| CFLAR | c-FLIP | - | Negative regulator of apoptosis and is structurally like caspase-8. | No | 51 | 0.59 |
| GPX4 | GPX4 | - | Oxidoreductase involved in ferroptosis. | Yes | 36 | 0.82 |
| SLC2A1 | GLUT1 | - | Glucose transporter. | Yes | 51 | 0.67 |
| TP63 | p63 | Oncogene or TSG | Transcription factor regulates stratified epithelial growth and homeostasis. | Yes | 48 | 0.6 |
| PKN2 | PRK2 | - | Serine/threonine kinase required for cell cycle progression and cell migration. | No | 36 | 0.58 |
| MAP3K11 | MLK3 | - | Serine/threonine kinase involved in apoptosis and cell migration. | No | 25 | 0.61 |
| TIPARP | PARP7 | Not Specified | ADP-ribose transferase involved in innate immunity and microtubule stability. | Yes | 38 | 0.68 |

Given the propensity of mutations and copy number variations to drive HNSCC, we investigated common genetic alterations to identify significant associations with the targetable dependencies from our pipeline. This led us to identify novel associations of *FAT1* alteration with both increased MBTPS1 and *TMX2* dependencies, which were both in our final list of 13 highest priority genes, and *NOTCH1* mutation with increased *TAP1* dependency. No known biologic connections could be made between these associations. Confirmatory *in-vitro* studies are necessary to further evaluate target efficacy in these molecular subgroups. These findings demonstrate the ability of our pipeline to enrich for dependencies found in only a subset of HPV(-) HNSCC cell line models. Identification of genetic subgroups that offer selective vulnerabilities may help improve therapeutic potential in future studies using the targets proposed in this pipeline.

A limitation of using cell lines is that they do not consider components of the tumor microenvironment during the *in vitro* screening process. Nonetheless, our study used DepMap based CRISPR-Cas9 screen and provided a systematic unbiased framework that effectively ranks oncogenic targets for further therapeutic targets development.

## CONCLUSION

The DepMap genome-wide loss of function CRISPR screens offer new insight into gene dependencies in cancer that not been previously identified. In this report, we present a strategy for prioritizing cancer-specific vulnerabilities most likely to offer a useful therapeutic window in cell line models. When applied to HPV(-) HNSCC, DepMap CRISPR screens capture well-studied targets in HNSCC in addition to 13 novel targets which had fitness effects as strong as the oncogene *PIK3CA*. We also demonstrate associations between the strength of these dependencies and the presence of common genetic alterations.Collectively, this study provides a catalog of gene dependencies that may be explored as potential therapeutic targets in HPV(-) HNSCC and a target prioritization strategy which can be readily applied to other cancer types.

## Notes

**Conflict of interest statement:** The authors have declared that no conflict of interest exists.

### Competing Interest Statement

The authors have declared no competing interest.

