## Supplemental Table for "Cancer type-specific prioritization strategy for targetable dependencies developed from the DepMap profile of head and neck cancer"

### SUPPLEMENTARY FIGURES AND TABLES

### INDEX

**Supplemental Table 1**: 143 targetable dependencies in cell line models of HNSCC

**Supplemental Table 2**: Functions of genes with clinical inhibitors in HNSCC and highest median gene effect scores

**Supplemental Figure 3**: Association between median gene effect score and copy number alterations or mutations in cell line models of HNSCC

### Supplemental Table 1: 143 targetable dependencies in cell line models of HNSCC.

| % Cell lines with essentiality | Median gene effect score | Gene symbol | Gene name | Known oncogene in OncoKB | Known inhibitor in DGIdb |
|---|---|---|---|---|---|
| 74 | 0.76 | MBTPS1 | membrane bound transcription factor peptidase, site 1 | No | Yes |
| 71 | 0.54 | KRT18 | keratin 18 | No | No |
| 70 | 0.71 | CDK6 | cyclin dependent kinase 6 | Yes | Yes |
| 68 | 0.59 | FARS2 | phenylalanyl-tRNA synthetase 2, mitochondrial | No | No |
| 66 | 0.6 | EGFR | epidermal growth factor receptor | Yes | Yes |
| 60 | 0.57 | PIK3CA | phosphatidylinositol-4,5-bisphosphate 3-kinase catalytic subunit alpha | Yes | Yes |
| 59 | 0.89 | TYMS | thymidylate synthetase | No | Yes |
| 57 | 0.56 | KLF5 | Kruppel like factor 5 | Yes | No |
| 56 | 0.68 | TUBB4B | tubulin beta 4B class IVb | No | Yes |
| 54 | 0.55 | UBA6 | ubiquitin like modifier activating enzyme 6 | No | No |
| 53 | 0.54 | RBM10 | RNA binding motif protein 10 | No | No |
| 52 | 0.55 | CARM1 | coactivator associated arginine methyltransferase 1 | No | Yes |
| 52 | 0.53 | NRDC | nardilysin convertase | No | No |
| 51 | 0.67 | SLC2A1 | solute carrier family 2 member 1 | No | No |
| 51 | 0.59 | CFLAR | CASP8 and FADD like apoptosis regulator | No | No |
| 51 | 0.52 | SSR2 | signal sequence receptor subunit 2 | No | No |
| 48 | 0.6 | TP63 | tumor protein p63 | Yes | No |
| 48 | 0.56 | ITGB1 | integrin subunit beta 1 | No | Yes |
| 47 | 0.49 | EARS2 | glutamyl-tRNA synthetase 2, mitochondrial | No | No |
| 46 | 0.47 | HNRNPA1 | heterogeneous nuclear ribonucleoprotein A1 | No | No |
| 45 | 0.59 | ERBB2 | erb-b2 receptor tyrosine kinase 2 | Yes | Yes |
| 43 | 0.56 | PPIH | peptidylprolyl isomerase H | No | No |
| 43 | 0.52 | HSD17B12 | hydroxysteroid 17-beta dehydrogenase 12 | No | No |
| 43 | 0.48 | SYT15 | synaptotagmin 15 | No | No |
| 41 | 0.55 | ERBB3 | erb-b2 receptor tyrosine kinase 3 | Yes | Yes |
| 41 | 0.48 | ASH2L | ASH2 like, histone lysine methyltransferase complex subunit | No | No |
| 41 | 0.47 | PMS2 | PMS1 homolog 2, mismatch repair system component | No | No |
| 40 | 0.67 | DHRSX | dehydrogenase/reductase X-linked | No | No |
| 40 | 0.53 | PTBP1 | polypyrimidine tract binding protein 1 | No | No |

| | | | | | |
|---|---|---|---|---|---|
| 40 | 0.48 | SLC3A2 | solute carrier family 3 member 2 | No | No |
| 39 | 0.51 | ODR4 | odr-4 GPCR localization factor homolog | No | No |
| 38 | 0.68 | TIPARP | TCDD inducible poly(ADP-ribose) polymerase | No | No |
| 38 | 0.49 | EXTL2 | exostosin like glycosyltransferase 2 | No | No |
| 37 | 0.58 | TMX2 | thioredoxin related transmembrane protein 2 | No | No |
| 37 | 0.54 | SLC4A7 | solute carrier family 4 member 7 | No | No |
| 37 | 0.49 | PTP4A1 | protein tyrosine phosphatase 4A1 | No | No |
| 37 | 0.47 | ZRANB2 | zinc finger RANBP2-type containing 2 | No | No |
| 36 | 0.82 | GPX4 | glutathione peroxidase 4 | No | No |
| 36 | 0.58 | PKN2 | protein kinase N2 | No | Yes |
| 34 | 0.5 | NDUFS3 | NADH:ubiquinone oxidoreductase core subunit S3 | No | Yes |
| 34 | 0.48 | SLC4A5 | solute carrier family 4 member 5 | No | No |
| 33 | 0.49 | CHMP1A | charged multivesicular body protein 1A | No | No |
| 32 | 0.53 | PPME1 | protein phosphatase methylesterase 1 | No | Yes |
| 31 | 0.47 | NISCH | nischarin | No | No |
| 30 | 0.49 | BOLA3 | bolA family member 3 | No | No |
| 30 | 0.45 | KCNJ9 | potassium inwardly rectifying channel subfamily J member 9 | No | Yes |
| 29 | 0.55 | MARK2 | microtubule affinity regulating kinase 2 | No | Yes |
| 29 | 0.5 | BIRC2 | baculoviral IAP repeat containing 2 | No | Yes |
| 28 | 0.54 | SPNS1 | sphingolipid transporter 1 (putative) | No | No |
| 28 | 0.46 | CSF3 | colony stimulating factor 3 | No | No |
| 26 | 0.51 | KCNA10 | potassium voltage-gated channel subfamily A member 10 | No | No |
| 26 | 0.49 | TXNDC9 | thioredoxin domain containing 9 | No | No |
| 26 | 0.49 | LCN1 | lipocalin 1 | No | No |
| 26 | 0.45 | ADAM11 | ADAM metallopeptidase domain 11 | No | No |
| 25 | 0.61 | MAP3K11 | mitogen-activated protein kinase kinase kinase 11 | No | Yes |
| 25 | 0.5 | RIPK3 | receptor interacting serine/threonine kinase 3 | No | Yes |
| 25 | 0.44 | ABCA4 | ATP binding cassette subfamily A member 4 | No | No |
| 25 | 0.43 | ENDOG | endonuclease G | No | No |
| 25 | 0.43 | INS | insulin | No | No |
| 24 | 0.52 | PAK2 | p21 (RAC1) activated kinase 2 | No | Yes |
| 24 | 0.47 | PROCA1 | protein interacting with cyclin A1 | No | No |
| 24 | 0.46 | H3C4 | H3 clustered histone 4 | No | No |
| 23 | 0.51 | CYB5R4 | cytochrome b5 reductase 4 | No | No |

| | | | | | |
|---|---|---|---|---|---|
| 23 | 0.49 | TYRO3 | TYRO3 protein tyrosine kinase | No | Yes |
| 23 | 0.48 | CNTNAP3B | contactin associated protein family member 3B | No | No |
| 22 | 0.62 | RAB35 | RAB35, member RAS oncogene family | Yes | No |
| 22 | 0.49 | IMPDH1 | inosine monophosphate dehydrogenase 1 | No | Yes |
| 22 | 0.49 | ELAVL1 | ELAV like RNA binding protein 1 | No | No |
| 22 | 0.48 | RNF31 | ring finger protein 31 | No | No |
| 22 | 0.48 | PI4KB | phosphatidylinositol 4-kinase beta | No | Yes |
| 22 | 0.48 | GRK2 | G protein-coupled receptor kinase 2 | No | Yes |
| 20 | 0.56 | PDE12 | phosphodiesterase 12 | No | No |
| 20 | 0.52 | ITGA3 | integrin subunit alpha 3 | No | No |
| 20 | 0.46 | PTAR1 | protein prenyltransferase alpha subunit repeat containing 1 | No | No |
| 20 | 0.45 | FASN | fatty acid synthase | No | Yes |
| 20 | 0.45 | ARTN | artemin | No | No |
| 18 | 0.47 | KCNK7 | potassium two pore domain channel subfamily K member 7 | No | No |
| 17 | 0.48 | ADAMTSL4 | ADAMTS like 4 | No | No |
| 17 | 0.47 | CYP4F11 | cytochrome P450 family 4 subfamily F member 11 | No | No |
| 17 | 0.46 | SCGB2A1 | secretoglobin family 2A member 1 | No | No |
| 17 | 0.45 | SLC29A2 | solute carrier family 29 member 2 | No | No |
| 17 | 0.43 | LTF | lactotransferrin | No | No |
| 16 | 0.62 | TDP2 | tyrosyl-DNA phosphodiesterase 2 | No | No |
| 16 | 0.47 | SLC25A33 | solute carrier family 25 member 33 | No | No |
| 16 | 0.46 | FGF19 | fibroblast growth factor 19 | Yes | No |
| 16 | 0.45 | SPINT1 | serine peptidase inhibitor, Kunitz type 1 | No | No |
| 16 | 0.43 | H4C5 | H4 clustered histone 5 | No | No |
| 16 | 0.39 | TAP1 | transporter 1, ATP binding cassette subfamily B member | No | No |
| 15 | 0.49 | EP300 | E1A binding protein p300 | No | Yes |
| 15 | 0.49 | UGCG | UDP-glucose ceramide glucosyltransferase | No | Yes |
| 15 | 0.46 | LDLR | low density lipoprotein receptor | No | No |
| 15 | 0.45 | MAP4K2 | mitogen-activated protein kinase kinase kinase kinase 2 | No | Yes |
| 15 | 0.44 | MANF | mesencephalic astrocyte derived neurotrophic factor | No | No |
| 15 | 0.44 | SLC25A1 | solute carrier family 25 member 1 | No | Yes |
| 15 | 0.38 | CYP2A13 | cytochrome P450 family 2 subfamily A member 13 | No | No |
| 14 | 0.54 | IGF1R | insulin like growth factor 1 receptor | Yes | Yes |
| 14 | 0.52 | SLC7A1 | solute carrier family 7 member 1 | No | No |

| | | | | | |
|---|---|---|---|---|---|
| 14 | 0.5 | CHST8 | carbohydrate sulfotransferase 8 | No | No |
| 14 | 0.48 | LMNA | lamin A/C | No | No |
| 14 | 0.48 | OVGP1 | oviductal glycoprotein 1 | No | No |
| 14 | 0.47 | GUCA2A | guanylate cyclase activator 2A | No | No |
| 14 | 0.46 | RPS6KB1 | ribosomal protein S6 kinase B1 | No | Yes |
| 14 | 0.46 | HCRTR1 | hypocretin receptor 1 | No | Yes |
| 14 | 0.45 | LGALS9 | galectin 9 | No | No |
| 14 | 0.45 | LIG1 | DNA ligase 1 | No | Yes |
| 14 | 0.42 | ITM2B | integral membrane protein 2B | No | No |
| 13 | 0.52 | TM2D1 | TM2 domain containing 1 | No | No |
| 13 | 0.51 | NAGLU | N-acetyl-alpha-glucosaminidase | No | No |
| 13 | 0.48 | HTRA2 | HtrA serine peptidase 2 | No | No |
| 13 | 0.46 | CDK8 | cyclin dependent kinase 8 | Yes | Yes |
| 13 | 0.46 | EIF4H | eukaryotic translation initiation factor 4H | No | No |
| 13 | 0.43 | HNRNPUL2 | heterogeneous nuclear ribonucleoprotein U like 2 | No | No |
| 13 | 0.43 | ABCB10 | ATP binding cassette subfamily B member 10 | No | No |
| 13 | 0.39 | ADAMTS7 | ADAM metallopeptidase with thrombospondin type 1 motif 7 | No | No |
| 11 | 0.53 | LTB4R2 | leukotriene B4 receptor 2 | No | Yes |
| 11 | 0.5 | P2RY6 | pyrimidinergic receptor P2Y6 | No | Yes |
| 11 | 0.49 | AHCYL1 | adenosylhomocysteinase like 1 | No | No |
| 11 | 0.49 | RNF123 | ring finger protein 123 | No | No |
| 11 | 0.48 | PCSK9 | proprotein convertase subtilisin/kexin type 9 | No | Yes |
| 11 | 0.47 | RBM5 | RNA binding motif protein 5 | No | No |
| 11 | 0.47 | LDHA | lactate dehydrogenase A | No | Yes |
| 11 | 0.45 | HAO2 | hydroxyacid oxidase 2 | No | No |
| 11 | 0.44 | SLC22A25 | solute carrier family 22 member 25 | No | No |
| 11 | 0.43 | LTBP3 | latent transforming growth factor beta binding protein 3 | No | No |
| 11 | 0.41 | PTPN1 | protein tyrosine phosphatase non-receptor type 1 | Yes | Yes |
| 10 | 0.52 | GSTM3 | glutathione S-transferase mu 3 | No | No |
| 10 | 0.5 | OGA | O-GlcNAcase | No | No |
| 10 | 0.45 | RTN3 | reticulon 3 | No | No |
| 10 | 0.45 | GANAB | glucosidase II alpha subunit | No | Yes |
| 10 | 0.44 | RTN4IP1 | reticulon 4 interacting protein 1 | No | No |
| 10 | 0.43 | GGTLC1 | gamma-glutamyltransferase light chain 1 | No | No |

| | | | | | |
|---|---|---|---|---|---|
| 10 | 0.41 | TNFRSF8 | TNF receptor superfamily member 8 | No | Yes |
| 9 | 0.51 | EGLN1 | egl-9 family hypoxia inducible factor 1 | No | Yes |
| 9 | 0.46 | PRKACA | protein kinase cAMP-activated catalytic subunit alpha | Yes | Yes |
| 9 | 0.46 | ASIC1 | acid sensing ion channel subunit 1 | No | No |
| 9 | 0.46 | CCL7 | C-C motif chemokine ligand 7 | No | No |
| 9 | 0.46 | TAC4 | tachykinin precursor 4 | No | No |
| 9 | 0.45 | CDC25B | cell division cycle 25B | No | No |
| 9 | 0.43 | MRGPRX3 | MAS related GPR family member X3 | No | No |
| 9 | 0.42 | CCS | copper chaperone for superoxide dismutase | No | No |
| 9 | 0.42 | RPS6KA4 | ribosomal protein S6 kinase A4 | Yes | Yes |
| 9 | 0.41 | MTCH2 | mitochondrial carrier 2 | No | No |
| 9 | 0.39 | SLC25A25 | solute carrier family 25 member 25 | No | No |

**Supplemental Table 2: Functions of genes with clinical inhibitors in HNSCC and highest median gene effect scores.**

| Gene symbol | Protein name | Designation in OncoKB | Description | Mechanistic study in HNSCC? | % Cell lines with essentiality | Median gene effect score |
|---|---|---|---|---|---|---|
| *CDK6* | CDK6 | Oncogene | Serine/threonine kinase that drives cell cycle progression. | Yes | 70 | 0.71 |
| *EGFR* | EGFR | Oncogene | Tyrosine kinase involved in growth signalling. | Yes | 66 | 0.6 |
| *PIK3CA* | p110α | Oncogene | Kinase catalytic subunit of PI3K, which regulates cell survival and proliferation. | Yes | 60 | 0.57 |
| *TUBB4B* | tubulin beta-4B | - | Microtubule subunit. | Yes | 56 | 0.68 |
| *TYMS* | TS | - | Enzyme in DNA synthesis. | Yes | 59 | 0.89 |
| *ERBB2* | HER2 | Oncogene | Tyrosine kinase involved in growth signalling. | Yes | 45 | 0.59 |

**Supplemental Figure 3**

**(A)**

| Gene name | Putative role | % with mutation in TCGA | % with CNA in TCGA | # Cell lines with CNA | Included in analysis? | # Cell lines with mutation | Mutations also included in analysis? | Total # cell lines included |
|---|---|---|---|---|---|---|---|---|
| PIK3CA | oncogene | 18.4 | 20.9 | 21 | Yes | 7 | Yes | 28 |
| CDKN2A | tumor suppressor | 22.1 | 30.3 | 21 | Yes | 13 | Yes | 34 |
| NOTCH1 | tumor suppressor | 17.8 | 2.1 | 7 | No | 4 | - | - |
| FAT1 | tumor suppressor | 22.5 | 7.3 | 31 | Yes | 10 | Yes | 41 |
| TP53 | mixed | 71.5 | 1.4 | 3 | No | 41 | - | - |
| EGFR | oncogene | 3.7 | 10.7 | 26 | Yes | 3 | No | 29 |

**(B)**

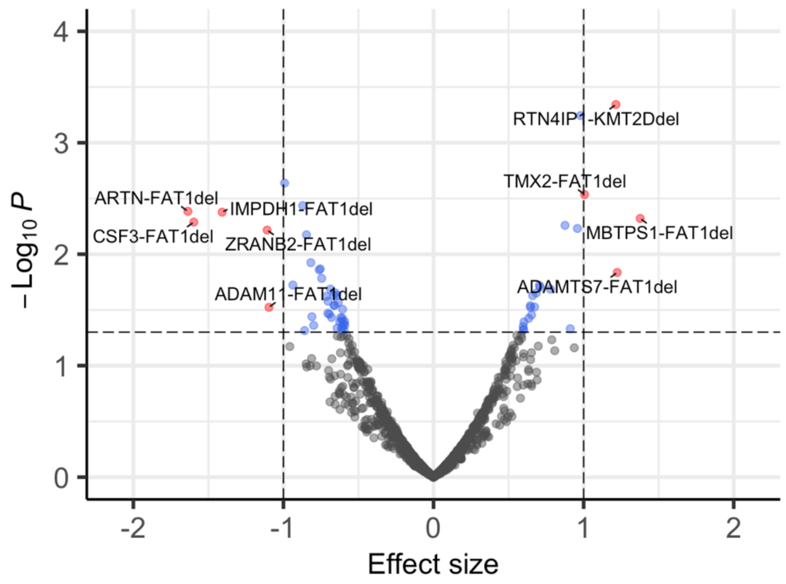


**Supplemental Figure 3: Association between median gene effect score and copy number alterations or mutations in cell line models of HNSCC :** (A) Known copy number alterations and mutations, and (B) Comparison with median gene effect score, with significant associations highlighted in red. A positive effect size represents increased dependency when the alteration is present.
